## Supplemental Figures and Tables for "Cellular underpinnings of the selective vulnerability to tauopathic insults in Alzheimer’s disease"

**Figure S1: Correlation structure of the Yao, *et al.* cell types. A.** Heat map of the Pearson correlations of the gene expression profiles of the Yao, *et al.* [1] cell types. **B.** Heat map of the Pearson correlations of the regional distributions of the Yao, *et al.* cell types as inferred by MISS [2].

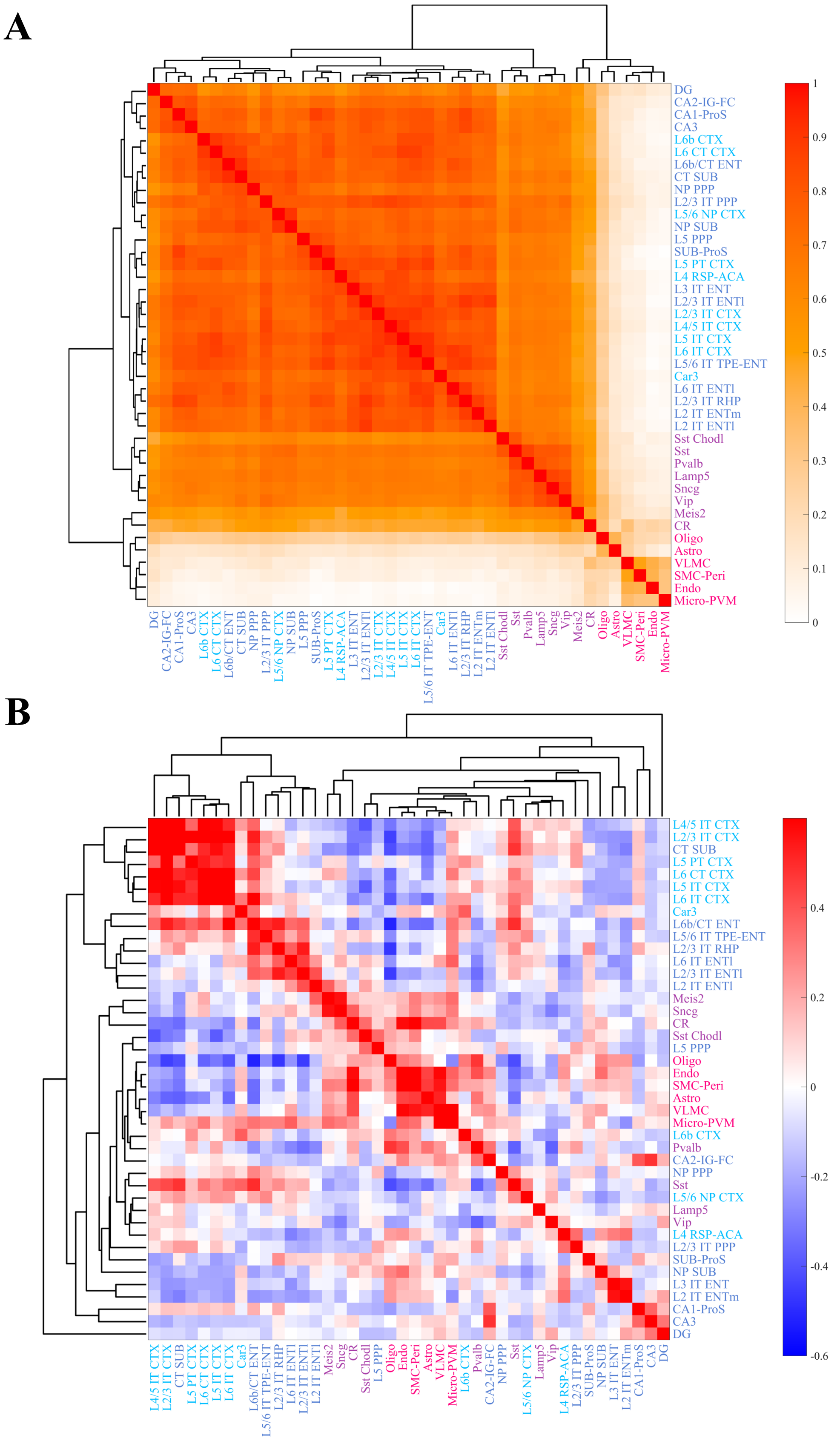
A
DG
CA2-IG-FC
CA1-ProS
CA3
L6b CTX
L6 CT CTX
L6b/CT ENT
CT SUB
NP PPP
L2/3 IT PPP
L5/6 NP CTX
NP SUB
L5 PPP
SUB-ProS
L5 PT CTX
L4 RSP-ACA
L3 IT ENT
L2/3 IT ENTl
L2/3 IT CTX
L4/5 IT CTX
L5 IT CTX
L6 IT CTX
L5/6 IT TPE-ENT
Car3
L6 IT ENTl
L2/3 IT RHP
L2 IT ENTm
L2 IT ENTl
Sst Chodl
Sst
Pvalb
Lamp5
Sncg
Vip
Meis2
CR
Oligo
Astro
VLMC
SMC-Peri
Endo
Micro-PVM
1
0.9
0.8
0.7
0.6
0.5
0.4
0.3
0.2
0.1
0
B
L4/5 IT CTX
L2/3 IT CTX
CT SUB
L5 PT CTX
L6 CT CTX
L5 IT CTX
L6 IT CTX
Car3
L6b/CT ENT
L5/6 IT TPE-ENT
L2/3 IT RHP
L6 IT ENTl
L2/3 IT ENTl
L2 IT ENTl
Meis2
Sncg
CR
Sst Chodl
L5 PPP
Oligo
Endo
SMC-Peri
Astro
VLMC
Micro-PVM
L6b CTX
Pvalb
CA2-IG-FC
NP PPP
Sst
L5/6 NP CTX
Lamp5
Vip
L4 RSP-ACA
L2/3 IT PPP
SUB-ProS
NP SUB
L3 IT ENT
L2 IT ENTm
CA1-ProS
CA3
DG
0.4
0.2
0
-0.2
-0.4
-0.6

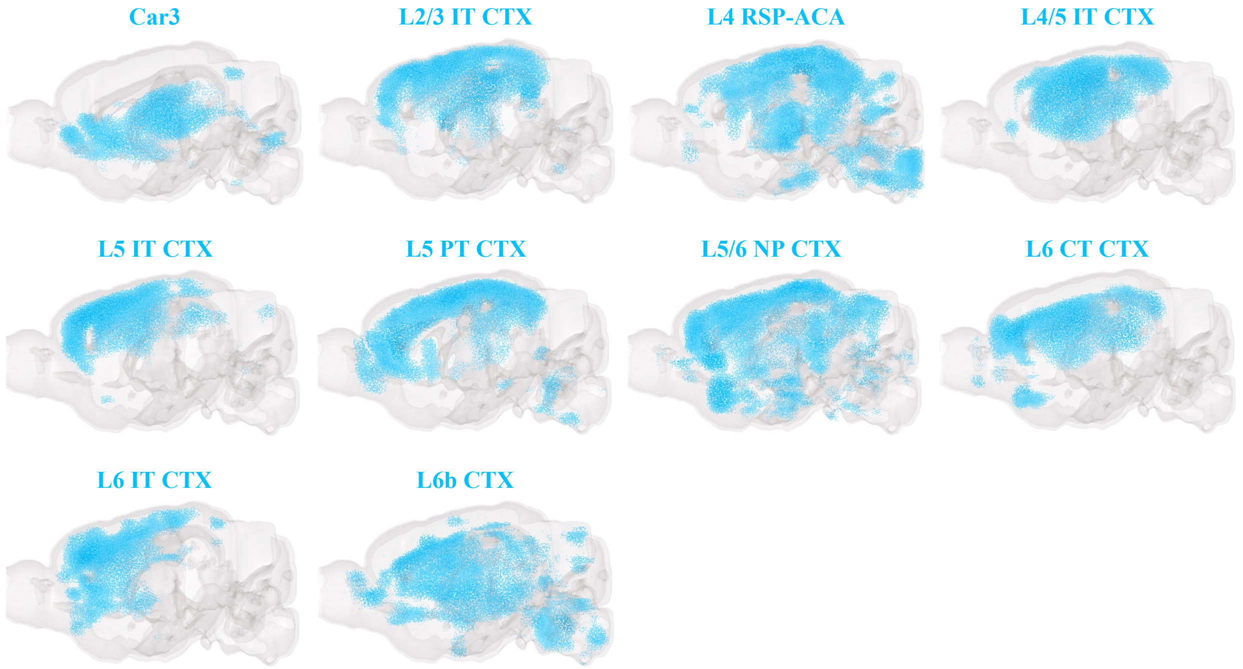


**Figure S2: Distributions of cortical glutamatergic neurons.** Sagittal views of the three-dimensional reconstructions of brain-wide densities of the cortical glutamatergic neurons in the Yao, *et al.* dataset [1]. Refer to **Table S1** and the original manuscript for further details on these cell types.

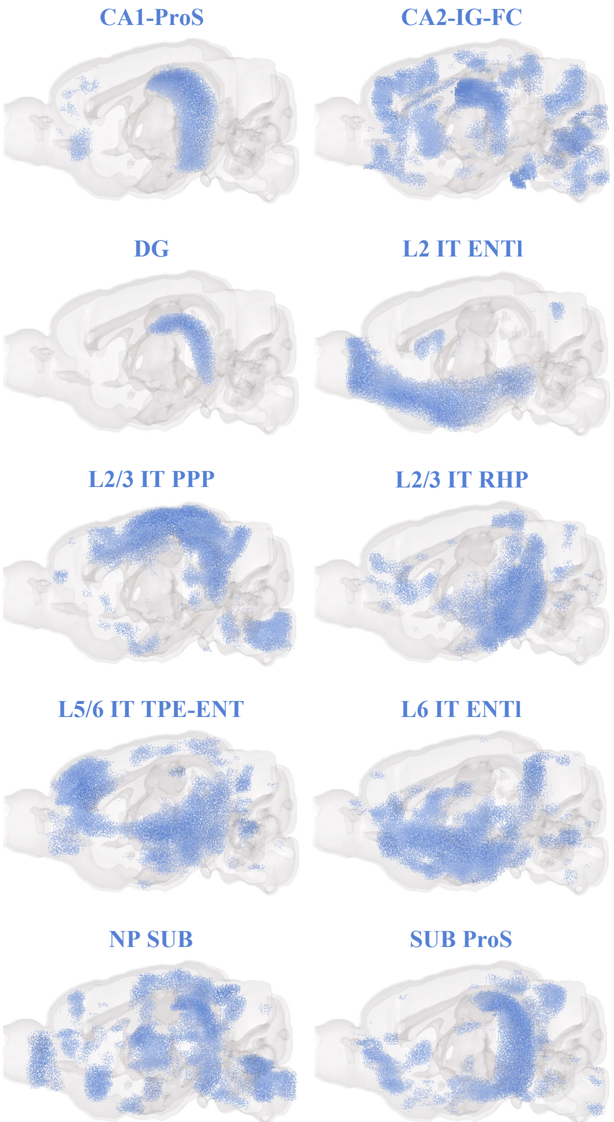


**Figure S3: Distributions of hippocampal glutamatergic neurons.** Sagittal views of the three-dimensional reconstructions of brain-wide densities of the hippocampal glutamatergic neurons in the Yao, *et al.* dataset [1]. Refer to **Table S1** and the original manuscript for further details on these cell types.

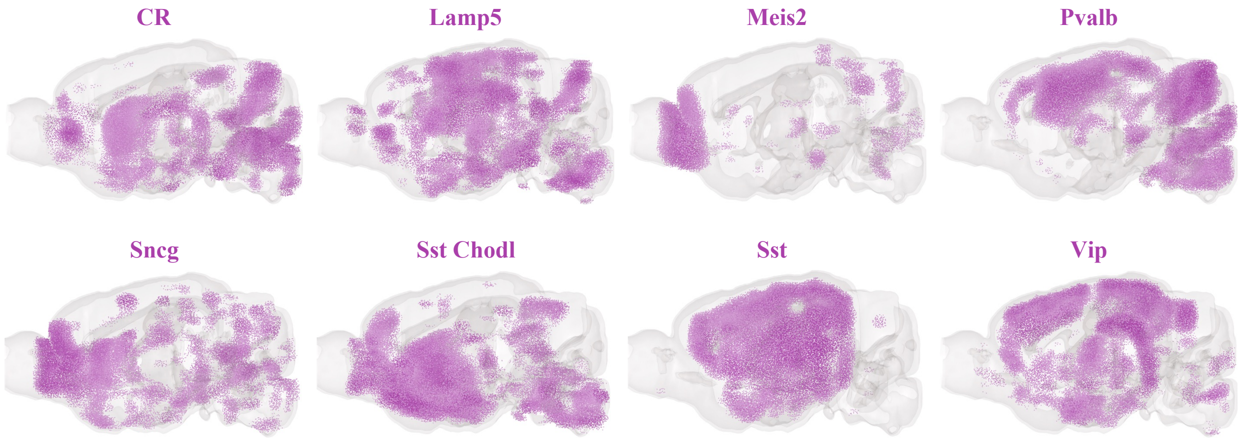


**Figure S4: Distributions of GABAergic neurons.** Sagittal views of the three-dimensional reconstructions of brain-wide densities of the GABAergic neurons in the Yao, *et al.* dataset [1]. Refer to **Table S2** and the original manuscript for further details on these cell types.

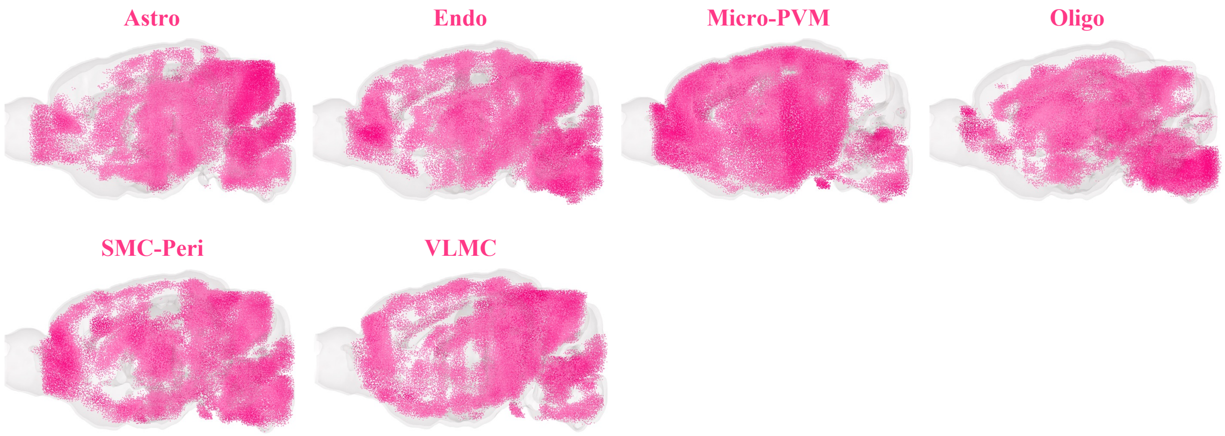


**Figure S5: Distributions of non-neuronal cells.** Sagittal views of the three-dimensional reconstructions of brain-wide densities of the non-neuronal cell types in the Yao, *et al.* dataset [1]. Refer to **Table S2** and the original manuscript for further details on these cell types.

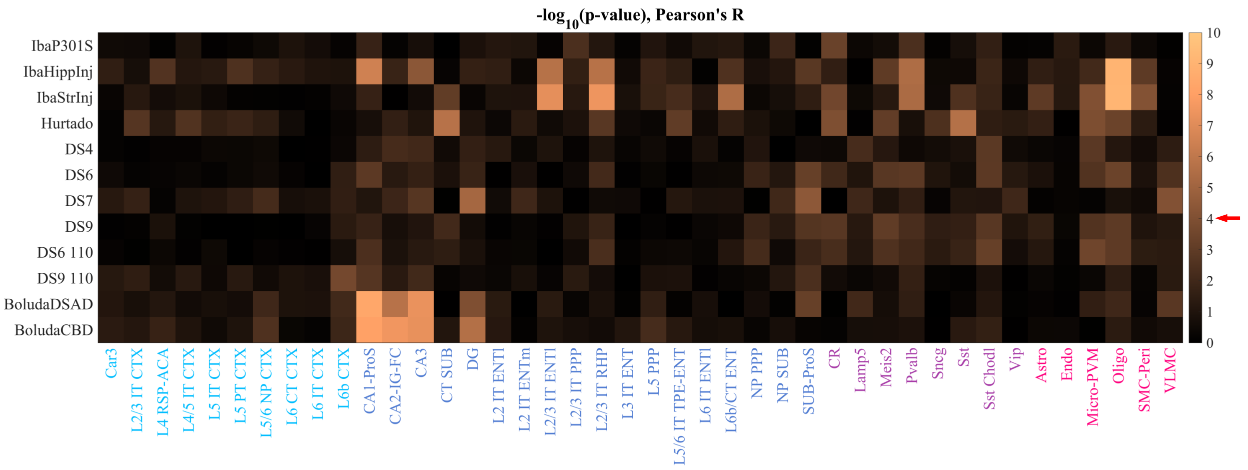


**Figure S6: Statistical significance of the correlations in Figure 2.** Heat map of the nominal -$\log_{10}$(p) values for the correlations presented in **Figure 2A**. The critical value corresponding to a Bonferroni-corrected significance level of 0.05 is -4.0 (red arrow).

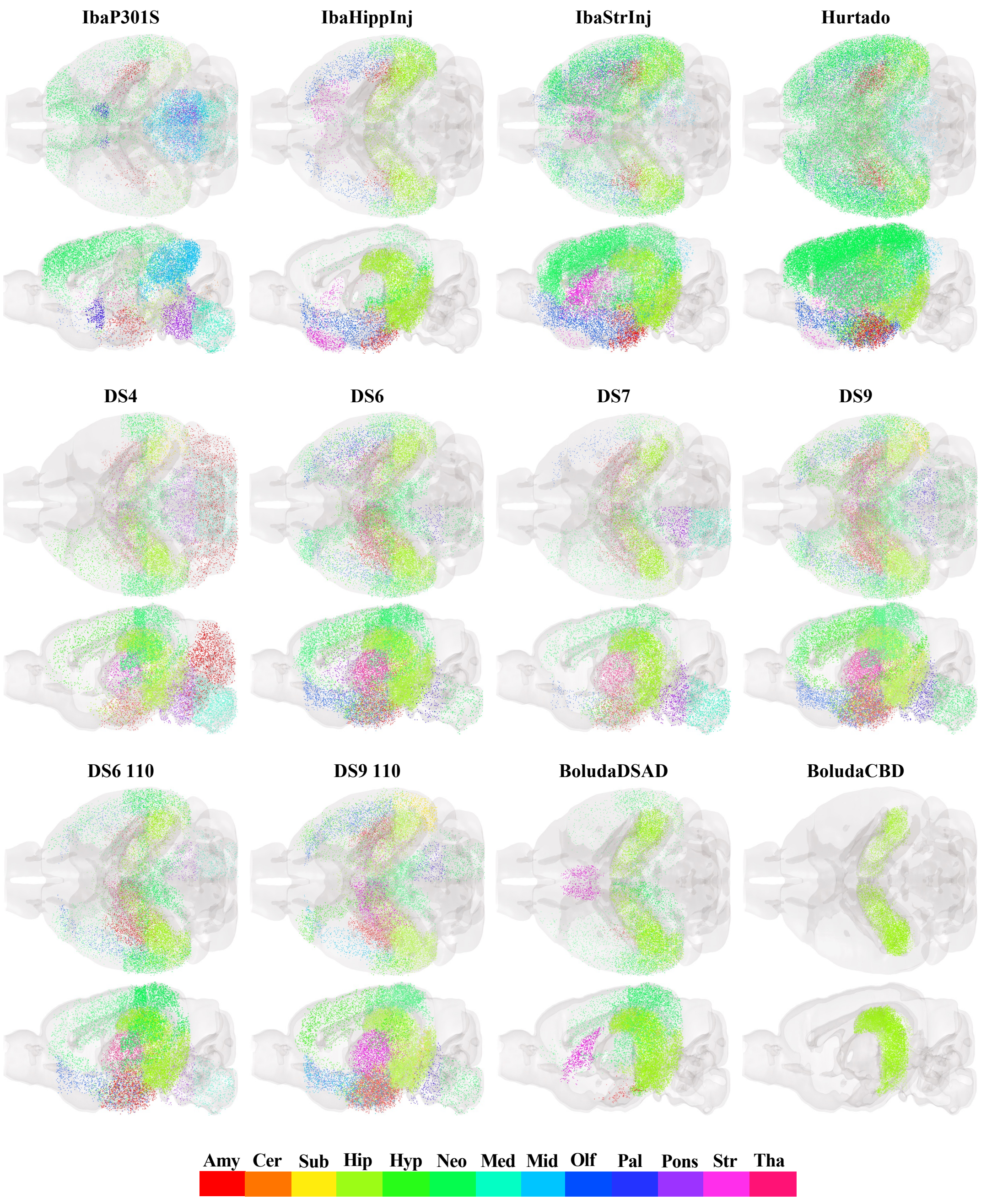


**Figure S7: End-timepoint pathology glass brains.** End-timepoint pathology for each of the twelve mouse tauopathy datasets, plotted in axial and sagittal views. See **Table S3** for descriptions of these datasets.

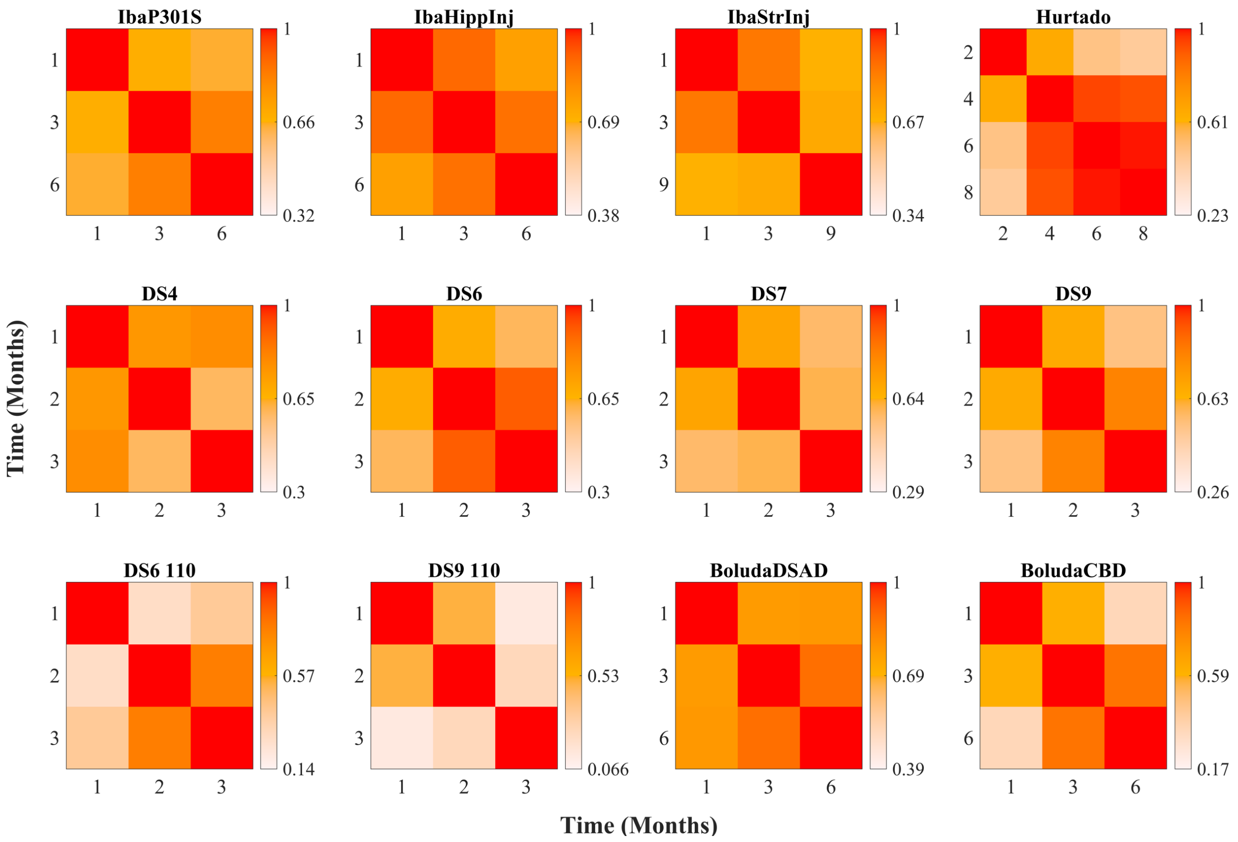


**Figure S8: Correlation structure of the mouse tauopathy datasets** Heat maps of the Pearson correlations between time points of the nine mouse tauopathy experiments analyzed in this study [3, 4, 5, 6, 7].

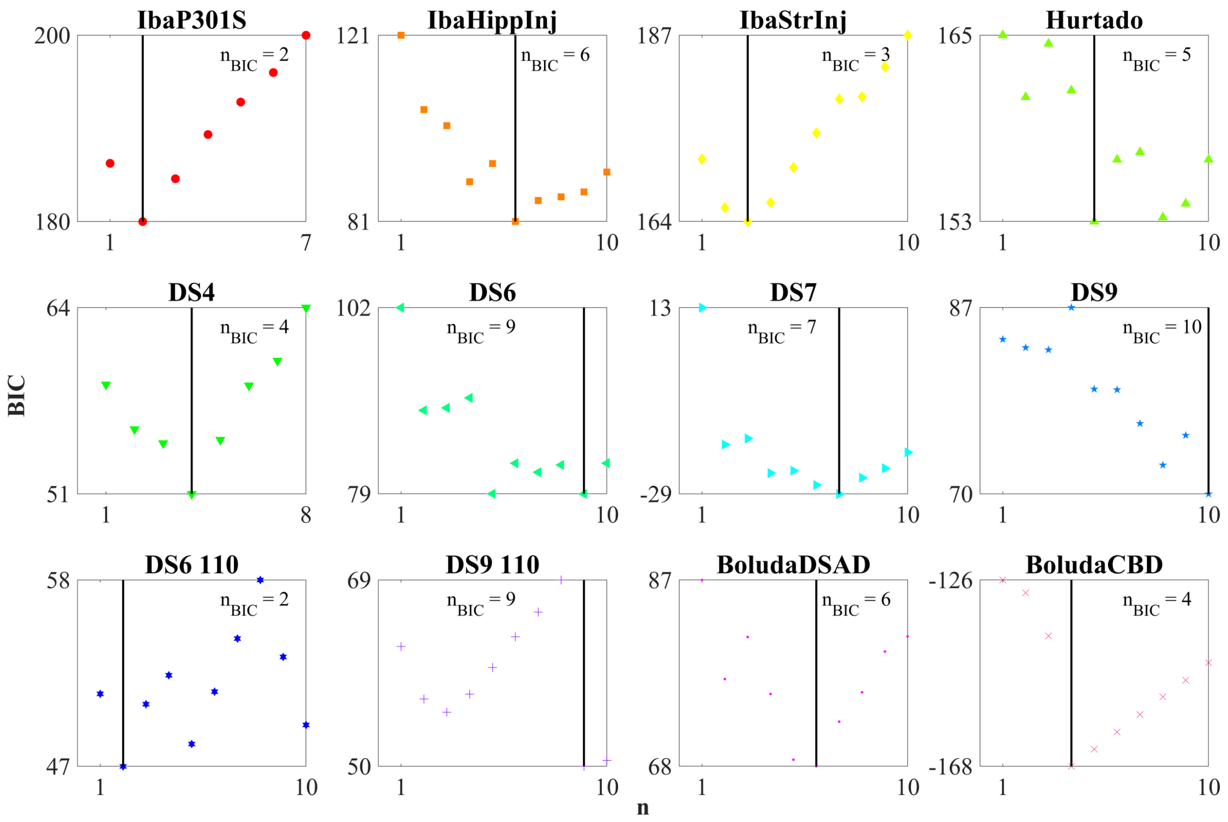


**Figure S9: BIC plots for the multivariate linear models in Figure 3.** Scatter plots of the BIC criterion with respect to the number of cell types added to the model (n) to determine the optimal sets for each tauopathy dataset.

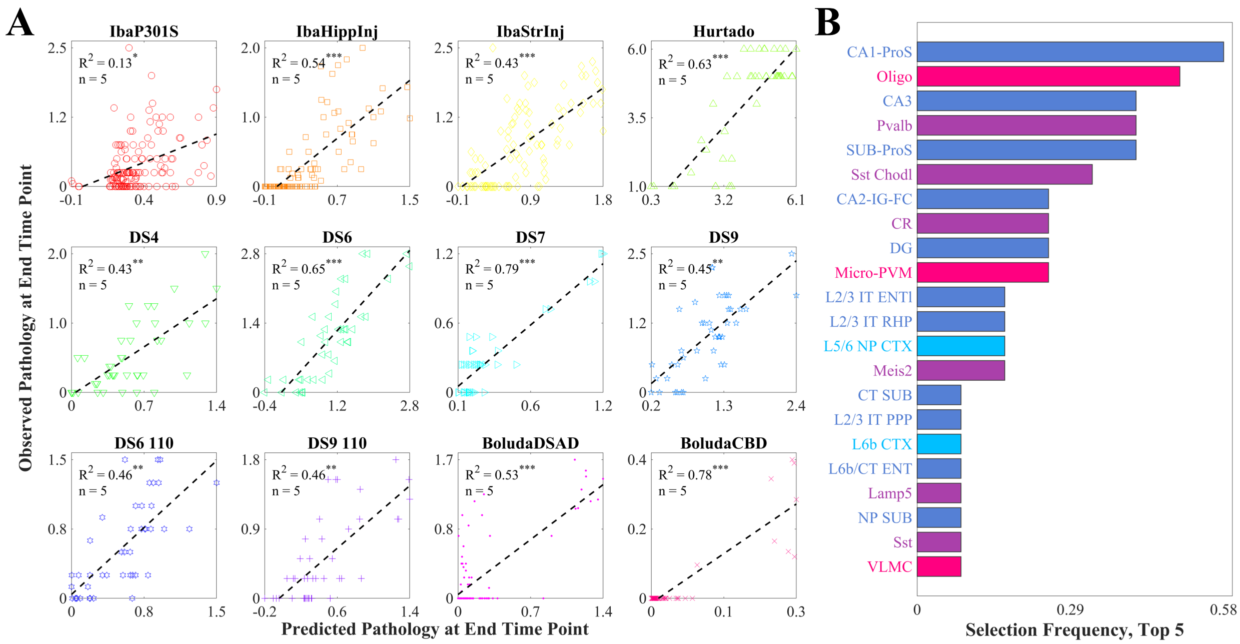


**Figure S10: Multivariate analysis of end-timepoint pathology, top five cell types A.** Scatter plots of the optimal cell-type-based models of tau pathology at the end time points for each of the nine mouse tauopathy studies, along with their associated $R^2$ values and the 5 cell types with the highest correlations to end-timepoint pathology (See **Figure 2**). **B.** Bar plot of the frequency with which cell types were included in the linear models in **A**. Of the 42 cell types in the Yao, *et al.* dataset, 22 were selected at least once.

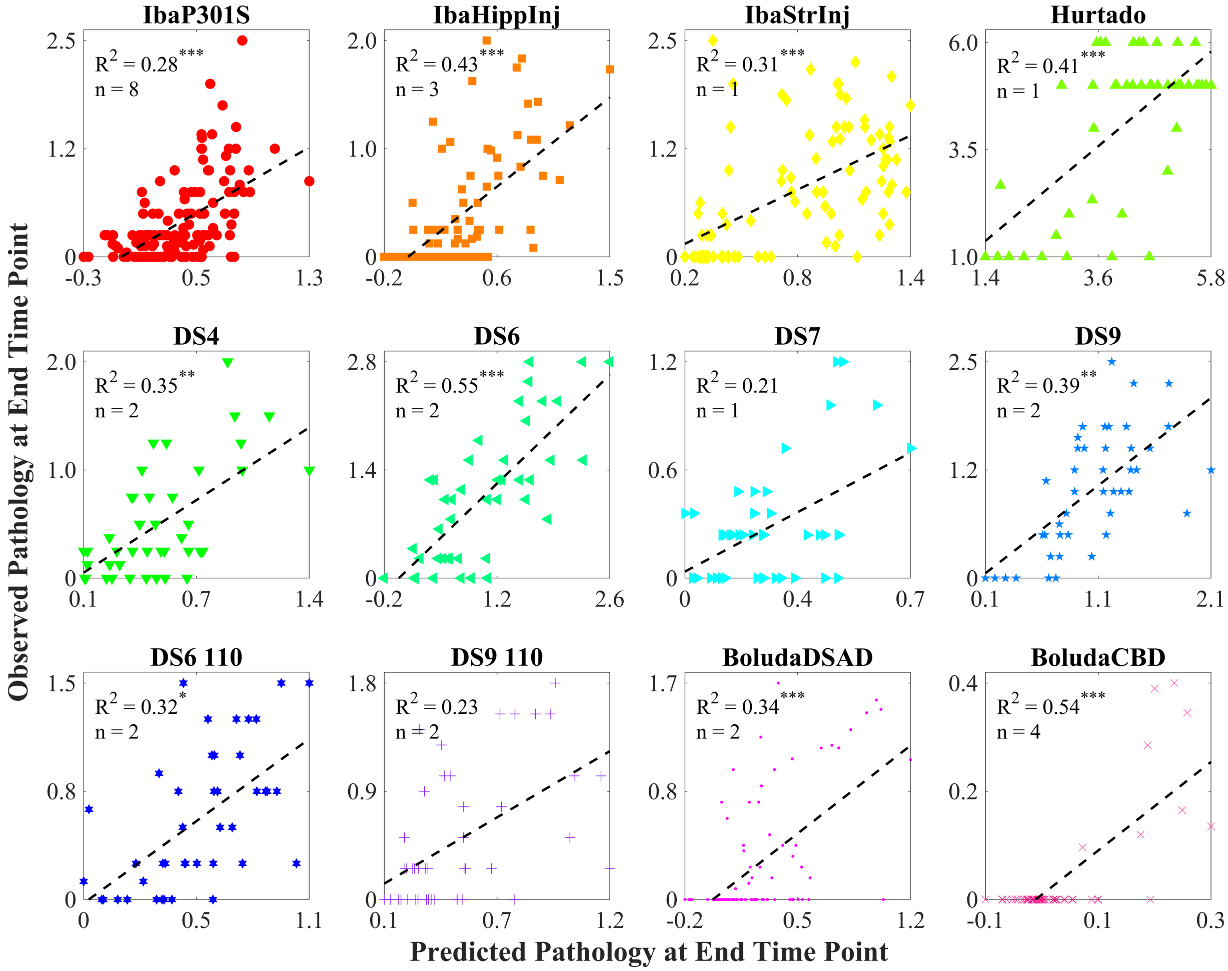


**Figure S11: Multivariate analysis of end-timepoint pathology, AD genes (BIC).** Scatter plots of the optimal cell-type-based models of tau pathology at the end time points for each of the nine mouse tauopathy studies, along with their associated $R^2$ values and the BIC-selected genes.

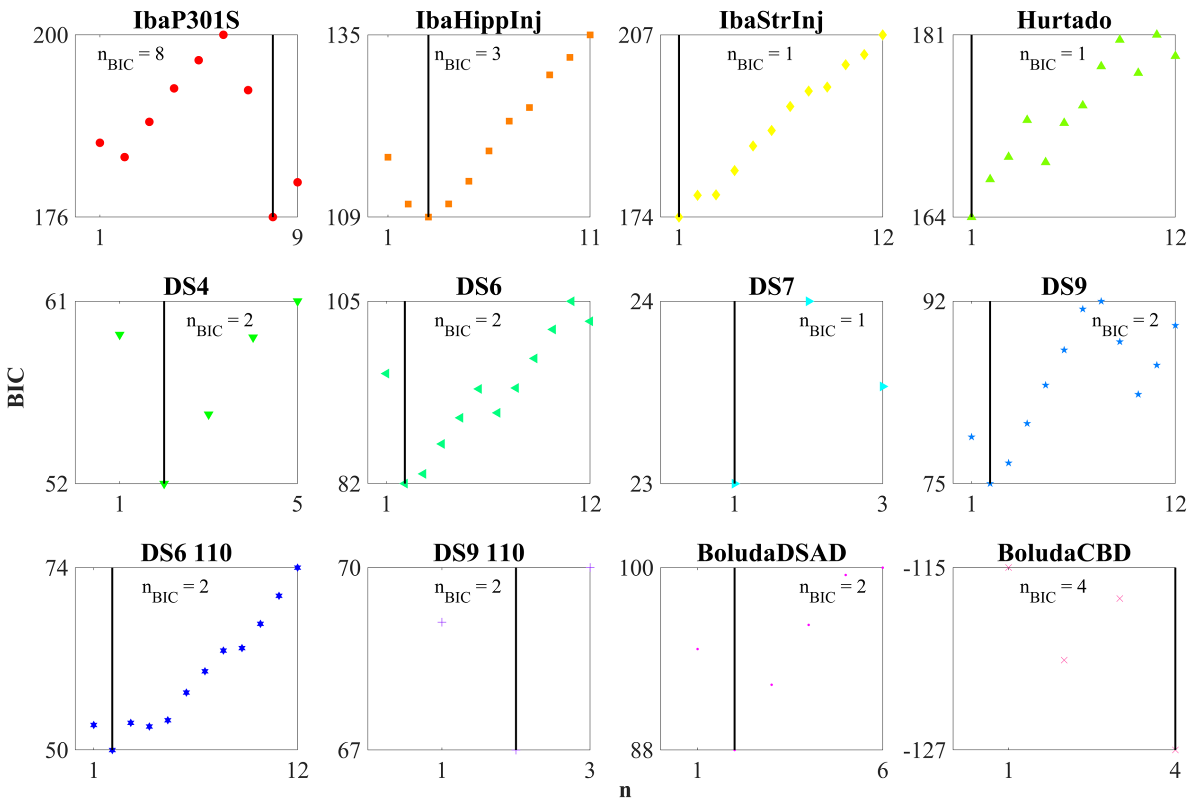


**Figure S12: BIC plots for the multivariate linear models in Figure S11.** Scatter plots of the BIC criterion with respect to the number of cell types added to the model (n) to determine the optimal sets for each tauopathy dataset.

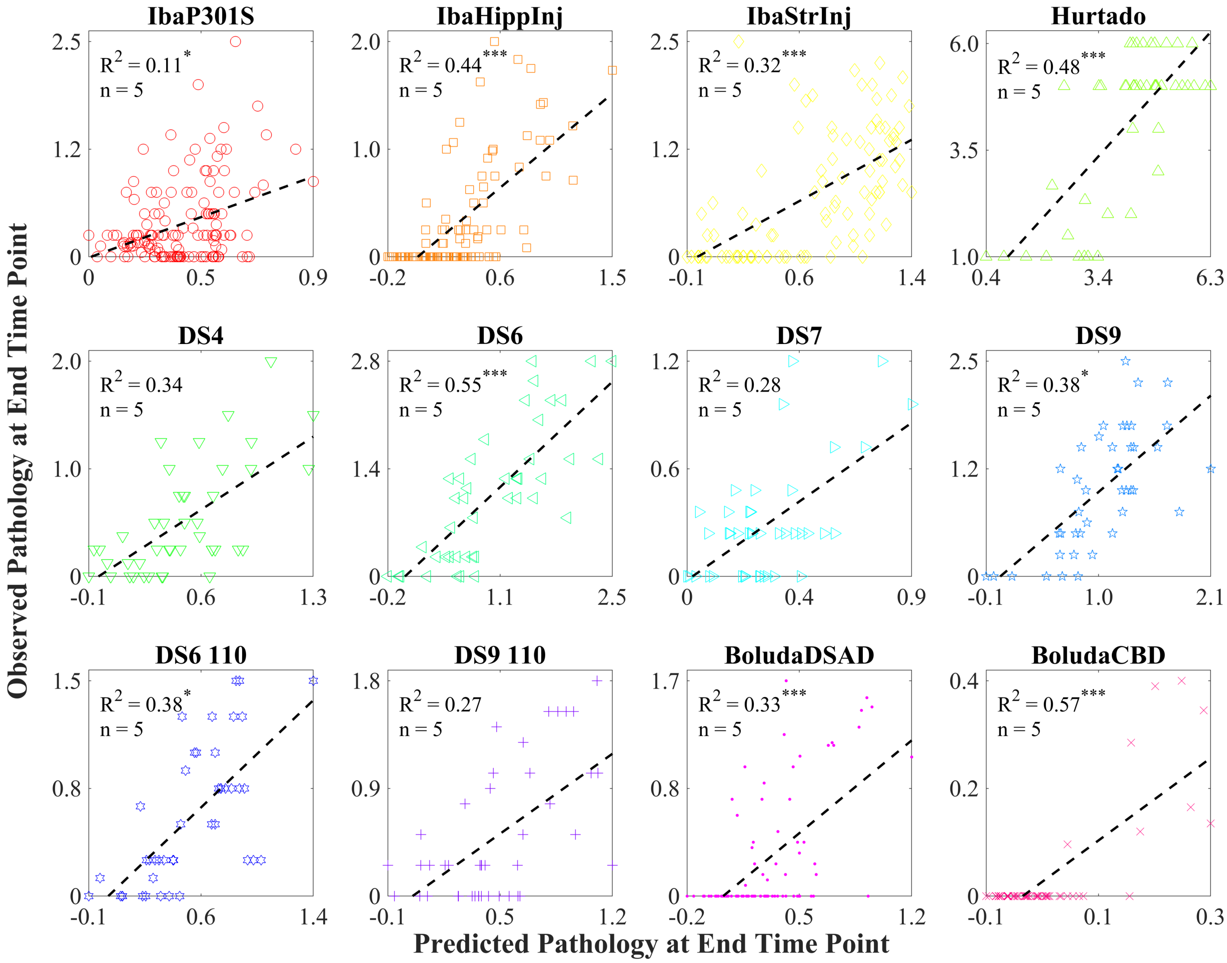


**Figure S13: Multivariate analysis of end-timepoint pathology, AD genes.** Scatter plots of the optimal cell-type-based models of tau pathology at the end time points for each of the nine mouse tauopathy studies, along with their associated $R^2$ values and the 5 genes with the highest correlations to pathology.

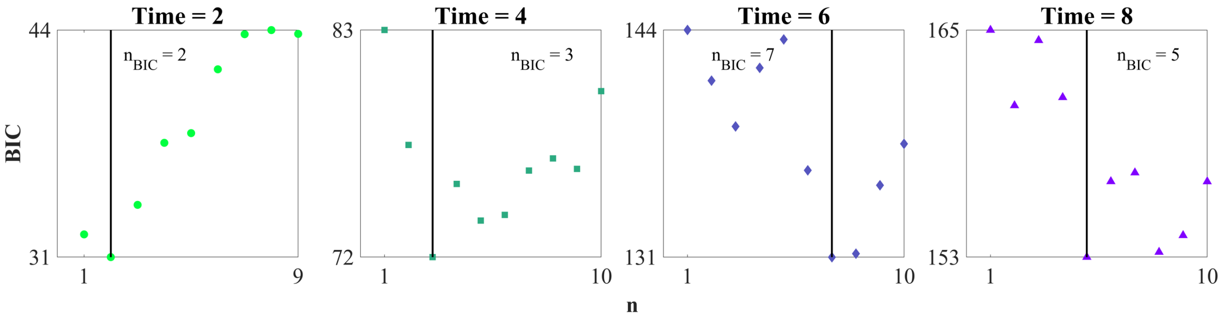


**Figure S14: BIC plots for the multivariate linear models in Figure 4.** Scatter plots of the BIC criterion with respect to the number of cell types added to the model (n) to determine the optimal sets the Hurtado, *et al.* dataset [4] for each timepoint.

### Supplemental Tables

| **Cortical glutamatergic neurons** | | **Hippocampal glutamatergic neurons** | |
|---|---|---|---|
| *Abbreviation* | *Full name* | *Abbreviation* | *Full name* |
| Car3 | *Car3*-expressing | CA1-ProS | CA1/prosubiculum |
| L2/3 IT CTX | Layer-2/3 intratelencephalic | CA2-FC-IG | CA2/fasciola cinereal/ induseum griseum |
| L4 RSP-ACA | Layer-4 retrospenial/ anterior cingulate | CA3 | CA3 |
| L4/5 IT CTX | Layer-4/5 intratelencephalic | CT SUB | Corticothalamic subiculum |
| L5 IT CTX | Layer-5 intratelencephalic | DG | Dentate gyrus |
| L5 PT CTX | Layer-5 pyramidal tract | L2 IT ENTl | Layer-2 intratelencephalic lateral entorhinal cortex |
| L5/6 NP CTX | Layer-5/6 near-projecting | L2 IT ENTm | Layer-2 intratelencephalic medial entorhinal cortex |
| L6 CT CTX | Layer-6 corticothalamic | L2/3 IT ENTl | Layer-2/3 intratelencephalic lateral entorhinal cortex |
| L6 IT CTX | Layer-6 intratelencephalic | L2/3 IT PPP | Layer-2/3 intratelencephalic para/post/presubiculum |
| L6b CTX | Layer-6b | L2/3 IT RHP | Layer-2/3 intratelencephalic retrohippocampal |
| | | L3 IT ENT | Layer-3 intratelencephalic entorhinal cortex |
| | | L5 PPP | Layer-5 para/post/presubiculum |
| | | L5/6 IT TPE-ENT | Layer-5/6 intratelencephalic temporal association/perirhinal/ ectorhinal/entorhinal |
| | | L6 IT ENTl | Layer-6 intratelencephalic lateral entorhinal cortex |
| | | L6b/CT ENT | Layer-6b/corticothalamic entorhinal cortex |
| | | NP PPP | Near-projecting para/post/ presubiculum |
| | | NP SUB | Near-projecting subiculum |
| | | SUB-ProS | Subiculum/prosubiculum |

**Table S1: Glutamatergic cell types**. List of the abbreviations and names of the glutamatergic cell types used in this study, each of which corresponds to a taxonomic subclass annotated by Yao *et al.* [1]. We have delineated these subclasses as being either cortical or hippocampal based on the annotations of their lower-level clusters.

| **GABAergic neurons** | | **Non-neuronal cells** | |
|---|---|---|---|
| *Abbreviation* | *Full name* | *Abbreviation* | *Full name* |
| CR | Cajal-Retzius | Astro | Astrocytes |
| Lamp5 | *Lamp5*-expressing | Endo | Endothelial |
| Meis2 | *Meis2*-expressing | Micro-PVM | Microglia/perivascular macrophages |
| Pvalb | *Pvalb*-expressing | Oligo | Oligodendrocytes |
| Sncg | *Sncg*-expressing | SMC-Peri | Smooth muscle cells/pericytes |
| Sst | *Sst*-expressing | VLMC | Vascular and leptomeningeal cells |
| Sst Chodl | *Sst*-expressing, long-range | | |
| Vip | *Vip*-expressing | | |

**Table S2: GABAergic and non-neuronal cell types**. List of the abbreviations and names of the GABAergic and non-neuronal cell types used in this study, each of which corresponds to a taxonomic subclass annotated by Yao *et al.* [1]. At the level of subclasses, these types are not uniquely defined between cortical and hippocampal regions.

| Name | Model | Injection site | Injectate | $n_{ROI}$ |
|---|---|---|---|---|
| IbaP301S [6] | PS19 | Locus coeruleus (RH) | Synthetic PFFs from 2N4R P301S τ (T40/PS) | 148 |
| IbaHippInj [5] | PS19 | Dentate gyrus (RH) | Synthetic PFFs from 2N4R P301S τ (T40/PS) and from truncated P301L τ (K18/PL) | 102 |
| IbaStrInj [5] | PS19 | Caudoputamen (RH) and primary motor area (RH) | Synthetic PFFs from 2N4R P301S τ (T40/PS) and from truncated P301L τ (K18/PL) | 96 |
| Hurtado [4] | PS19/ PDAPP | None | None | 45 |
| DS4 [7] | PS19 | CA1 (LH) | Isolated from AD brain homogenate; prominent nuclear inclusions ("speckles") | 44 |
| DS6 [7] | PS19 | CA1 (LH) | Isolated from P301S mouse brain homogenate; fibril-like cytoplasmic inclusions ("threads") | 44 |
| DS7 [7] | PS19 | CA1 (LH) | Recombinant fibrils; prominent nuclear inclusions ("speckles") | 44 |
| DS9 [7] | PS19 | CA1 (LH) | Recombinant fibrils; prominent nuclear inclusions ("speckles") | 44 |
| DS6 110 [7] | PS19 | CA1 (LH) | DS6 strain, 1:10 dilution | 44 |
| DS9 110 [7] | PS19 | CA1 (LH) | DS9 strain, 1:10 dilution | 44 |
| BoludaDSAD [3] | PS19 | CA1 (LH) and primary somatosensory area (LH) | DSAD brain homogenate | 90 |
| BoludaCBD [3] | PS19 | CA1 (LH) and primary somatosensory area (LH) | CBD brain homogenate | 58 |

**Table S3: Mouse tauopathy datasets**. List of the tauopathy datasets explored here with descriptions of four key experimental features: mouse genetic background, injection site, type of τ injected, and the number of regions for which τ pathology was quantified. All studies quantified τ pathology within hemispheres ipsilateral and contralateral to the injection site separately with the exception of Hurtado, which was bilaterally averaged. RH – right hemisphere; LH – left hemisphere; PFF – preformed fibrils; DSAD – Down Syndrome Alzheimer's disease; CBD – corticobasal dengeneration.

| **One-way t-test** | **t-statistic** | $p_{\text{MHC}}$ | **Two-way t-test** | **t-statistic** | $p_{\text{MHC}}$ |
|---|---|---|---|---|---|
| *Cortical glutamatergic* | -5.20 | $3.32 \times 10^{-6}$ | *Cortical glut. – Hippocampal glut.* | -7.72 | $8.07 \times 10^{-13}$ |
| *Hippocampal glut.* | 6.79 | $4.27 \times 10^{-10}$ | *Cortical glut. – GABAergic* | 0.320 | 1 |
| *GABAergic* | -3.32 | $5.09 \times 10^{-3}$ | *Cortical glut. – Non-neuronal* | -1.49 | 0.827 |
| *Non-neuronal* | -0.964 | 1 | *Hippocampal glut. – GABAergic* | 6.60 | $1.79 \times 10^{-10}$ |
| | | | *Hippocampal glut. – Non-neuronal* | 4.19 | $3.79 \times 10^{-5}$ |
| | | | *GABAergic – Non-neuronal* | -1.37 | 1 |

**Table S4: T-test results for cell-type classes**. Summary of one-way and two-way t-test results for distributions of Pearson's R values of the four cell-type classes within the Yao, *et al.* dataset: cortical glutamatergic neurons, hippocampal glutamatergic neurons, GABAergic neurons, and non-neuronal cells (see **Figure 2C**). T-tests were performed after first using the Fisher's R-to-Z transformation on the individual Pearson's R values displayed in **Figure 2A**. The $p$-values reported have been multiple-hypothesis corrected using the Bonferroni criterion. See **Tables S1** and **S2** for a complete list of the cell types within each class.

| Gene symbol | Gene name | Biological function |
|---|---|---|
| *Adamts1* | ADAM metallopeptidase with thrombospondin type 1 motif 1 | Extracellular matrix organization |
| *Ank3* | Ankyrin-3 | Membrane-cytoskeleton linker |
| *Apoe* | Apolipoprotein E | Negative regulation of apoptotic process |
| *App* | Amyloid-beta precursor protein | Axonogenesis, neurite growth, neuronal adhesion |
| *Bace1* | Beta-secretase 1 | Proteolysis of amyloid-beta precursor protein |
| *Cd33* | Myeloid cell surface antigen CD33 | Cell adhesion, cell-cell interactions |
| *Clu* | Clusterin | Extracellular chaperone protein |
| *Doc2a* | Double C2-like domain-containing protein alpha | $Ca^{2+}$-dependent neurotransmitter release |
| *Epdr1* | Mammalian ependymin-related protein 1 | Cell-matrix adhesion |
| *Grid2* | Glutamate receptor ionotropic, delta-2 | Glutamate receptor |
| *Grin2b* | Glutamate receptor ionotropic, NMDA 2B | Glutamate receptor |
| *Hs3st2* | Heparan sulfate glucosamine 3-O-sulfotransferase 2 | Glycosaminoglycan biosynthetic process |
| *Il34* | Interleukin-34 | Proliferation, survival and differentiation of monocytes and macrophages |
| *Mapk14* | Mitogen-activated protein kinase 14 | MAP kinase signalling pathway |
| *Mapt* | Microtubule-associated protein tau | Microtubule assembly and stabilization |
| *Pld3* | 5'-3' exonuclease PLD3 | Regulates inflammtory response to single-stranded DNA |
| *Prnp* | Major prion protein | Unclear primary biological function |
| *Rorb* | Nuclear receptor ROR-beta | DNA-binding transcription factor |
| *Sirpa* | Tyrosine-protein phosphatase non-receptor type substrate 1 | Cell surface receptor, cell adhesion |
| *Slc44a1* | Choline transporter-like protein 1 | Choline transporter |
| *Sorl1* | Protein Sortilin-related receptor | Intracellular protein trafficking and localization |
| *Spp1* | Osteopontin | Cell-matrix adhesion |
| *Tmem41a* | Transmembrane protein 41A | Unclear primary biological function |
| *Trem2* | Triggering receptor expressed on myeloid cells 2 | Disease-associated microglia activation |

**Table S5: AD-associated genes**. List of the names and brief descriptions of the genes examined using univariate (**Figure 4**) and multivariate (**Figure S11** and **Figure S13**) selective vulnerability analyses, each of which has one or more variants associated with AD incidence. These genes represent an intersection between the list given by the Alzheimer's Disease Sequencing Project [8, 9] and the coronal series of the Allen Gene Expression Atlas (AGEA) [10], which yielded 24 genes. Gene annotations were obtained from the UniProt database [11] unless otherwise noted.

| **Dataset** | **Cell types (top 5)** | | | **AD risk genes (top 5)** | | |
|---|---|---|---|---|---|---|
| | $R^2$ | n | BIC | $R^2$ | n | BIC |
| IbaP301S [6] | **0.13**$^{*}$ | **5** | **192.7** | 0.11$^{*}$ | 5 | 196.6 |
| IbaHippInj [5] | **0.54**$^{***}$ | **5** | **93.4** | 0.44$^{***}$ | 5 | 114.1 |
| IbaStrInj [5] | **0.43**$^{***}$ | **5** | **170.4** | 0.32$^{***}$ | 5 | 193.2 |
| Hurtado [4] | **0.64**$^{***}$ | **5** | **153.3** | 0.48$^{***}$ | 5 | 169.0 |
| DS4 [7] | **0.43**$^{**}$ | **5** | **54.5** | 0.34 | 5 | 61.2 |
| DS6 [7] | **0.65**$^{***}$ | **5** | **79.4** | 0.55$^{***}$ | 5 | 90.2 |
| DS7 [7] | **0.79**$^{***}$ | **5** | **-23.5** | 0.28 | 5 | 29.9 |
| DS9 [7] | **0.45**$^{**}$ | **5** | **79.4** | 0.38$^{*}$ | 5 | 84.3 |
| DS6 110 [7] | **0.46**$^{**}$ | **5** | **48.7** | 0.38$^{*}$ | 5 | 54.3 |
| DS9 110 [7] | **0.46**$^{**}$ | **5** | **60.0** | 0.27 | 5 | 72.8 |
| BoludaDSAD [3] | **0.53**$^{***}$ | **5** | **68.7** | 0.33$^{***}$ | 5 | 99.4 |
| BoludaCBD [3] | **0.78**$^{***}$ | **5** | **-163.9** | 0.57$^{***}$ | 5 | -128.1 |

**Table S6: Top-five feature linear model statistics**. Statistics corresponding to the linear models shown in **Figure S10** and **Figure S13**. Bold font indicates the best model by the Bayesian Information Criterion (BIC). *: $p < 0.01$; **: $p < 0.001$; ***: $p < 0.0001$.

| t (months) | $R^2$ | n | BIC | Most significant cell type | t-statistic | $p$-value |
|---|---|---|---|---|---|---|
| 2 | $0.28^{*}$ | 2 | 30.7 | L3 IT ENT | 2.9 | $6.7 \times 10^{-3}$ |
| 4 | $0.52^{***}$ | 3 | 71.2 | Sst | 3.7 | $6.2 \times 10^{-4}$ |
| 6 | $0.69^{***}$ | 7 | 131.2 | CT SUB | 3.3 | $2.1 \times 10^{-3}$ |
| 8 | $0.63^{***}$ | 5 | 153.3 | Oligo | -3.4 | $1.5 \times 10^{-3}$ |

**Table S7: Hurtado dataset linear model statistics**. Statistics corresponding to the linear models shown in **Figure 5A**, along with the cell type with the coefficient with the single-highest t-statistic. *: $p < 0.01$; **: $p < 0.001$; ***: $p < 0.0001$.

**Table S8-S11. Gene lists for SV-G, SV-C, SR-G, and SR-C gene ontology analysis.**
